# *PbrSYP71* regulates pollen tube growth by maintaining uneven distribution of the ER *in Pyrus*

**DOI:** 10.1101/2024.01.17.576111

**Authors:** Mingliang Zhang, Chao Tang, Chi Lan, Ningyi Zhang, Dong Yue, Zhihua Xie, Ming Qian, Mengjun Sun, Zongqi Liu, Zhu Xie, Hao Zhang, Zhuqin Liu, Shaoling Zhang, Peng Wang, Juyou Wu

**Author notes:** To whom correspondence should be addressed: Peng Wang,; Juyou Wu. These authors contributed equally (Mingliang Zhang, Chao Tang).

## Abstract

The uneven distribution of endoplasmic reticulum (ER) underlies the rapid polar growth of pollen tubes. However, the mechanism governing ER distribution remains elusive. In this study, we have identified a pollen tube-specific syntaxin protein, PbrSYP71. Our findings reveal that both overexpression and knocking down of *PbrSYP71* inhibited pollen tube growth. Subcellular localization analysis demonstrates that PbrSYP71 anchors to the ER via its transmembrane structure. Overexpression of *PbrSYP71* leads to clustered ER distribution in the pollen tube, while knocking down of *PbrSYP71* abolishes the uneven ER distribution. Remarkably, transient overexpression of *PbrSYP71*Δ*ABD*, lacking the actin binding domain (ABD) of PbrSYP71, has no impact on ER distribution or pollen tube growth. Further investigation indicates that ABD is positioned on F-actin in the pollen tube and has a direct interaction with F-actin. PbrSYP71 assists the ER in moving towards the apex of pollen tube, with ABD displaying autonomous mobility. Our study elucidates that PbrSYP71 maintains uneven distribution of the ER by tethering ER to F-actin, facilitating ER movement towards the pollen tube apex for pear pollen tube elongation. These insights shed light on the mechanisms governing ER distribution in polarized cell growth.

## Introduction

Pollen tube delivers male gametes to the ovaries for double fertilization, which is a necessary step for sexual reproduction in flowering plants. The polarized growth of pollen tube is a very fast process, which requires essential materials supported by secretory system consisting of the endoplasmic reticulum (ER), the Golgi apparatus, and vesicles (Steer and Steer, 1989; Campanoni and Blatt, 2007; Ruan *et al*., 2020). Meanwhile, pollen tube requires intense movements of secretory system to ensure the efficient delivery of these materials to the growing tip. These distribution and movements are not completely randomized, instead, specific classes of organelles and vesicles are aggregated at the pollen tube tip (Cai *et al*., 2015). For example, secretory vesicles and ER respectively accumulate at apical and subapical area of pollen tube (Parton *et al*., 2001; Lovy-Wheeler *et al*., 2007). Exploring the molecular mechanisms of secretion system distribution in pollen tube is crucial for understanding its rapid polarized growth.

The ER dynamically spreads throughout the cytoplasm and is crucial for cellular health (Westrate *et al*., 2015; Zheng *et al*., 2022). ER distribution is regulated by the actin filaments (F-actin), which is further mediated by actin-linker proteins. For now, two actin-linker proteins are identified, including AtSYP73 (a syntaxin protein of plants) and NET 3B in *Arabidopsis* (Cao *et al*., 2016; Wang and Hussey, 2017). The aggregation state of the ER in tobacco leaf is formed by an increase in the actin-linker protein and is decomposed by the depolymerization of the actin cytoskeleton (Cao *et al*., 2016; Wang and Hussey, 2017). In pollen tube, F-actin is found to regulate ER distribution (Lovy-Wheeler *et al*., 2007). The role of actin-linker proteins (i.e. AtSYP73 protein that is found to localized at ER) in regulating ER distribution in pollen tube, however, is still unknown.

AtSYP73 is a member of the plant SYP7 subgroup of the soluble N-ethylmaleimide-sensitive factor attachment protein receptor (SNARE) proteins, which is important for membrane fusion (Jahn and Scheller, 2006). SNAREs are divided into vesicle-associated SNAREs (v-SNAREs) and target cell-associated SNAREs (t-SNAREs) based on their subcellular localization, and are divided into R-SNARE and Q-SNARE based on their conserved residues. Q-SNARE is further classified into three types, i.e. Qa-, Qb-, and Qc-SNARE (Lipka *et al*., 2007). In general, R-SNAREs function as v-SNAREs, while Q-SNAREs act as t-SNAREs. v-SNAREs and t-SNAREs are two important components in exocytosis process. v-SNAREs are localized at vesicle, which need to be attached to “t-SNARE complex”, composed of three t-SNARE proteins localized at target membrane, to form a core four-helix SNARE bundle called trans-SNARE complex to finish the membrane fusion (Jahn and Scheller, 2006; Han *et al*., 2017). For tissues (e.g. leaves) in which ER is randomly distributed, exocytosis occurs at different directions, causing non-polarized growth of these tissues (Cao *et al*., 2016; Schwarz and Blower, 2016; Akita *et al*., 2017). In some tissues (e.g. pollen tube), ER is aggregated at their specific location (Campanoni and Blatt, 2007; Cheung and Wu, 2007; Lovy-Wheeler *et al*., 2007; Hepler and Winship, 2015), exocytosis thus potentially causes polarized growth of such tissues.

In “t-SNARE complex”, one helix always comes from syntaxin group (Weimbs *et al*., 1997; Sanderfoot *et al*., 2000). Syntaxin of plants (SYP) is categorized into eight subgroups (SYP1-8) as part of the Q-SNARE complex. Within *Arabidopsis* the SYP1, 2, 3, 4, and 8 subgroups are associated with Qa-SNARE, while the SYP5, 6, and 7 subgroups are linked to Qc-SNARE (Sanderfoot *et al*., 2000; Lipka *et al*., 2007). In SYP7 subgroup in *Arabidopsis*, AtSYP73 regulates the ER distribution and movement by interacting with actin in hypocotyl epidermal cells (Cao *et al*., 2016). It remains unclear whether the members of the SYP7 subgroup play a role in the ER distribution in the pollen tube.

Pear (*Pyrus bretschneideri*) is an important orchard crop worldwide, of which the pollination is a very laborious but necessary process due to its self-incompatibility. Breeding self-compatibility cultivars has always been an important goal in pear production, for which understanding the underlying mechanisms of pollen tube growth is very relevant. Here, we chose pear (*Pyrus bretschneideri* cv. ‘Dangshansuli’) as the case study due to its well-documented genome information that provides fundamental database for studying the molecular mechanisms of pollen tube growth (Wu *et al*., 2013). We identified a gene from the SYP7 subgroup, *PbrSYP71*, which was highly expressed in pear pollen tube and promoted pollen tube growth. We found that overexpression and knocking down of *PbrSYP71* both lead to inhibited pollen tube growth. Further research revealed that ER-located PbrSYP71 mediates the distribution of ER in the pollen tube, thereby affecting vesicle secretion. Next, we found that actin binding domain (ABD) is key domain for PbrSYP71 to regulating distribution of ER. Finally, we confirmed that PbrSYP71 binds to F-actin through ABD interacting with F-actin. Therefore, our results show how *PbrSYP71* regulates pollen tube growth by maintaining the uneven distribution of the ER.

## Results

### *PbrSYP* genes were identified in pear and *PbrSYP71* was specifically expressed in the pollen tube

A total of 43 *SYP* members from the pear genome were identified (*Table supplement 1*). These proteins were distributed in 8 subgroups (SYP1-SYP8) based on evolutionary relationships (*Figure 1—figure supplement 1*). The SYP7 subgroup included five members, i.e. *PbrSYP71*, *72*, *73*, *74*, and *75*, which had high level of homology and conservative domain and motif with *AtSYP71*, *72*, and *73* (*Figure 1A, B*; *Figure* 1—*figure supplement 2*, and *Figure* 1—*figure supplement 3*). Our RT-qPCR data showed that among all five *PbrSYPs*, only *PbrSYP71*, *73* and *75* were expressed in the pollen tube, with *PbrSYP71* showing the highest expression level (*Figure 1C*). Expressions of *PbrSYP72* and *74* were hardly detected in the pollen tube. Moreover, *PbrSYP71* was only highly expressed in the pollen tube but not in any other tissues e.g. leaves or stems (*Figure* 1—*figure supplement 4*, and *Figure* 1—*figure supplement 5*). These results suggested an important role of *PbrSYP71* in pear pollen tube growth.

**Figure 1.**
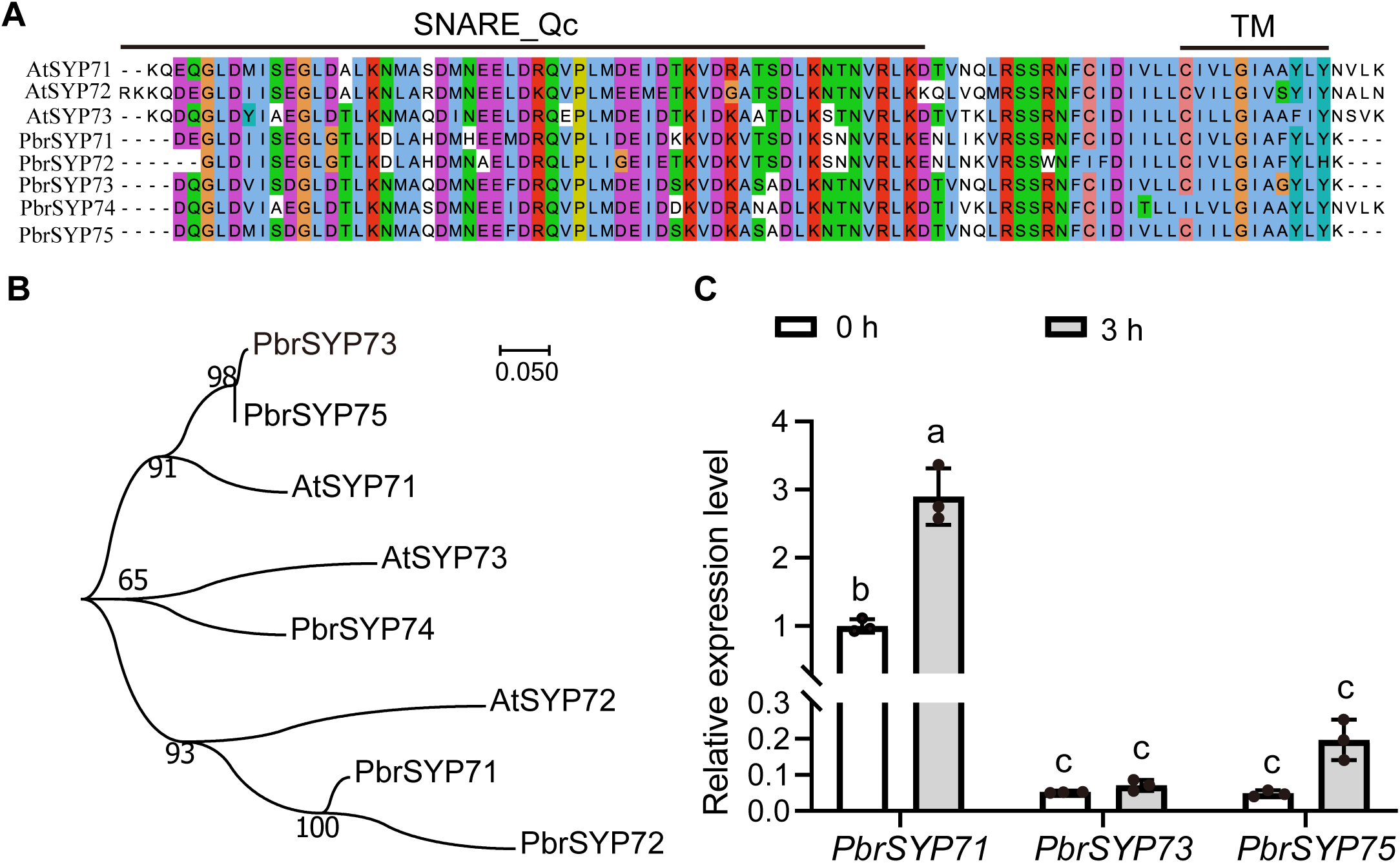
*PbrSYP71*, a member of the plant SYP7 subgroup of SNARE, was expressed in pear pollen tube. **A)** Sequence alignment of PbrSYP71, PbrSYP72, PbrSYP73, PbrSYP74, PbrSYP75, AtSYP71, AtSYP72 and AtSYP73. The black line represents the SNARE-Qc domain and transmembrane (TM) domain. **B)** Phylogenetic relationship of pear and *Arabidopsis* SYP7 subgroup proteins. The tree was generated using the Maximum Likelihood method in MEGA7. **C)** Expression level of *PbrSYP71*, *PbrSYP73*, and *PbrSYP75* in development of pollen tube (0 h and 3 h) were determined by RT-qPCR assay. Data are mean± SEM of 3 independent biological replicates. Different letters indicate significant differences as determined by ANOVA followed by Tukey’s multiple comparison test (*p* < 0.05).

### *PbrSYP71* regulated pear pollen tube growth

To verify the function of *PbrSYP71* in pear pollen tube growth, Firstly, we overexpressed *PbrSYP71-GFP* in pear pollen by particle bombardment, with *GFP* as control. Overexpression of *PbrSYP71* resulted in significantly lower pollen tube length compared to the control group (*Figure 2A, C*). This result indicated that overexpression of *PbrSYP71* inhibited pear pollen tube growth.

**Figure 2.**
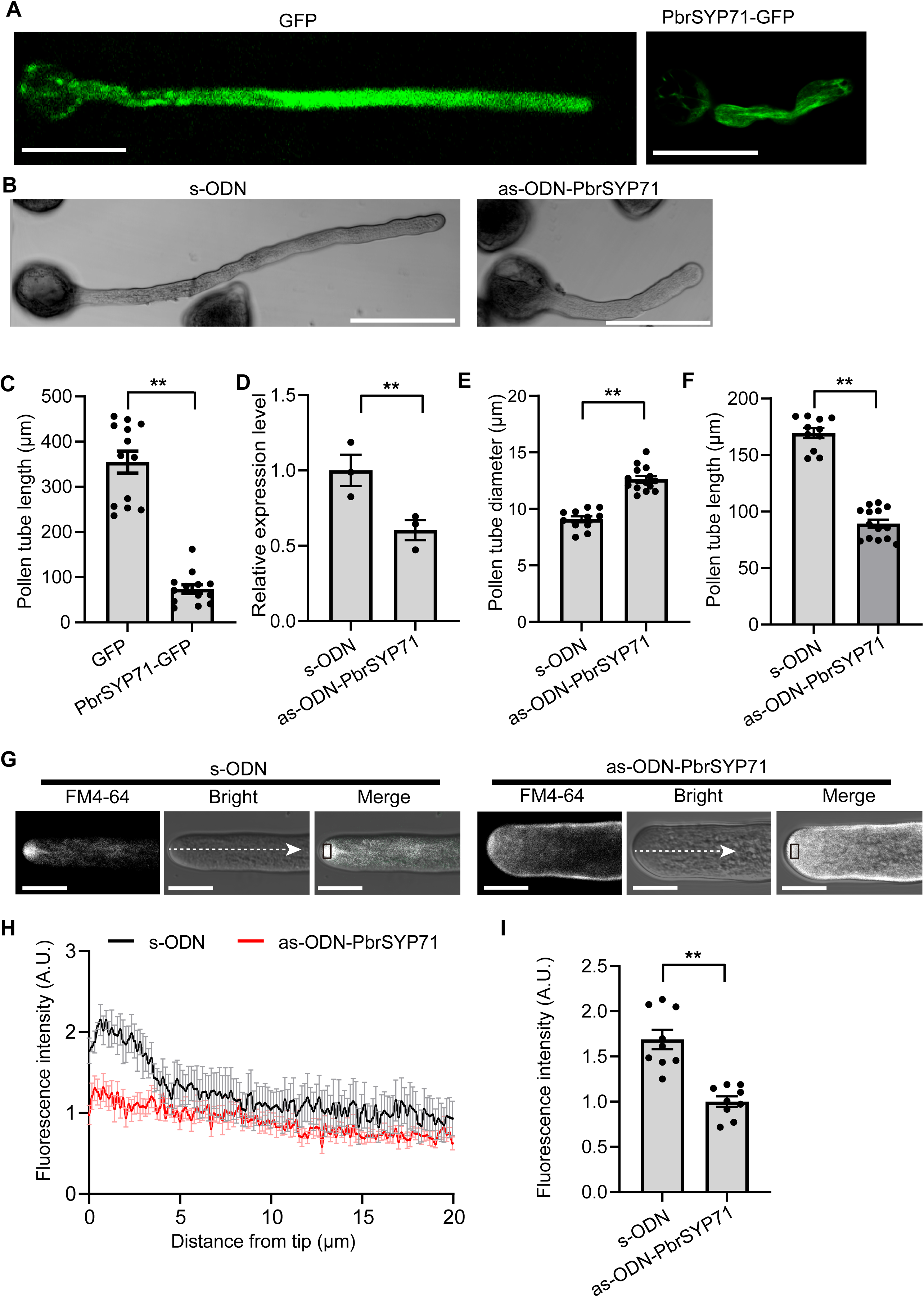
*PbrSYP71* regulates pear pollen tubes growth by affecting the content of secretory vesicles. **A)** Image of pear pollen tube transiently expressing GFP or PbrSYP71-GFP. The exprssion of *PbrSYP71-GFP* and *GFP* were driven by the NTP303 promoter. PbrSYP1-GFP and GFP were transiently expressed in pear pollen tubes by particle bombardment. The pollen tube was cultured for 6 h after transformation. Scale bars = 50 μm. **B)** The typical Image of pear pollen tube treated with sense oligodeoxynucleotide (s-ODN), as control, or antisense oligodeoxynucleotide of PbrSYP71 (as-ODN-PbrSYP71) for 3 h. Scale bars = 50 μm. **C)** Quantitative analysis of pollen tube length shown in (A), n = 13 pollen tubes. **D)** The relative exprssion level of *PbrSYP71* in pollen tube transfected with s-ODN, and as-ODN-PbrSYP71, determined by RT-qPCR assays, n = 3. **E and F)** Quantitative analysis of pollen tube diameter and length in (B), respectively, n≥ 9 pollen tubes. **G)** The typical Image of secretory vesicles of pear pollen tube treated with s-ODN or as-ODN-PbrSYP71, secretory vesicles were stained by FM4-64 dye, Scale bars = 10 μm. **H)** Fluorescence intensity of secretory vesicles at 20 µm distance from pollen tube tips along the white thread in (G), n = 5. **I)** Quantitative analysis of FM4-64 fluorescence intensity of pollen apcial area at black box in (G), n = 9. For (C-F) and (I), data are mean± SEM, differences were identified using Student’s *t*-test, significant differences between means are indicated by **(*p* < 0.01).

Meanwhile, the *PbrSYP71* expression was knocked down using antisense oligodeoxynucleotide (ODN) silencing technology with s-ODN as control. The phenotype of pear pollen tubes was mal-functioning when treated with as-ODN-PbrSYP71 (*Figure 2B*). The as-ODN-PbrSYP71 treatment significantly reduced *PbrSYP71* expression level (*Figure. 2D*), but did not affect the expression levels of *PbrSYP73* and *PbrSYP75* (*Figure* 2—*figure supplement 1*). The pollen tube length was significantly decreased whereas pollen tube diameter was increased in the presence of as-ODN-PbrSYP71 (*Figure 2E, F*). We further examined the content of secretory vesicles at the apex of pollen tubes using FM4-64 labeling. The content of secretory vesicles under treatment with as-ODN-PbrSYP71 was significantly lower than that in the control group (*Figure 2G-I*). In rapidly growing pollen tubes, secretory vesicles are always accumulated at the apex of pear pollen tube (*Movie supplement 1*). After knocking down the expression of *PbrSYP71*, the content of secretory vesicles at the apex of pollen tube significantly decreased, and the growth rate of pollen tubes slowed down significantly (*Movie supplement 2*). The cellulose content of the pollen tube was significantly decreased when the *PbrSYP71* expression was knocked down (*Figure* 2—*figure supplement 2*). These results suggested that knocking down the expression of *PbrSYP71* caused pollen tube growth defects by affecting the secretion vesicle content, which presented a similar outcome with the overexpression of *PbrSYP71*.

### The ER-localized PbrSYP71 reshaped the ER distribution in pollen tube

To analyze the subcellular localization of PbrSYP71, we transiently overexpressed *PbrSYP71* in pear pollen tubes and tobacco epidermal leaves, respectively. In pear pollen tube, NTP303: GFP and NTP303: mCherry were uniformly and continuously distributed in the cytoplasm of pollen tube cells (*Figure 3* — *figure supplement 1*). ER was marked by HDEL-mCherry using particle bombardment. HDEL-mCherry was distributed in regions other than the apex, with the most distribution in the subapical region and uniform distribution in the shank region (*Figure 3A-C*). After co-overexpression of PbrSYP71-GFP and HDEL-mCherry in pollen tubes, HDEL-mCherry showed a clustered distribution in pollen tube shank as shown by the white arrows in Figure 3D (*Figure 3D-F*). PbrSYP-GFP and HDEL showed similar distribution patterns, with a high pearson correlation coefficient of 0.828± 0.057 (*Figure 3D*). These results indicated that PbrSYP71 anchored to the ER, and overexpression of *PbrSYP71* affected the distribution of the ER. To further verify whether the transmembrane (TM) domain of the PbrSYP71 protein anchored to the ER, we constructed a PbrSYP71-TM-GFP vector containing the TM domain (236-265 aa) (*Figure 3*—f*igure supplement 2A*). Then, we co-overexpressed PbrSYP71-TM-GFP and HDEL-mCherry in the pollen tube, the distribution of HDEL-mCherry remained unchanged (*Figure 3G-I*). PbrSYP71-TM-GFP and HDEL-mCherry showed similar distributions, with a high pearson correlation coefficient of 0.824± 0.040 (*Figure 3H*). These results indicated that PbrSYP71-TM was located in ER, and overexpression of PbrSYP71-TM did not affect the distribution of ER.

**Figure 3.**
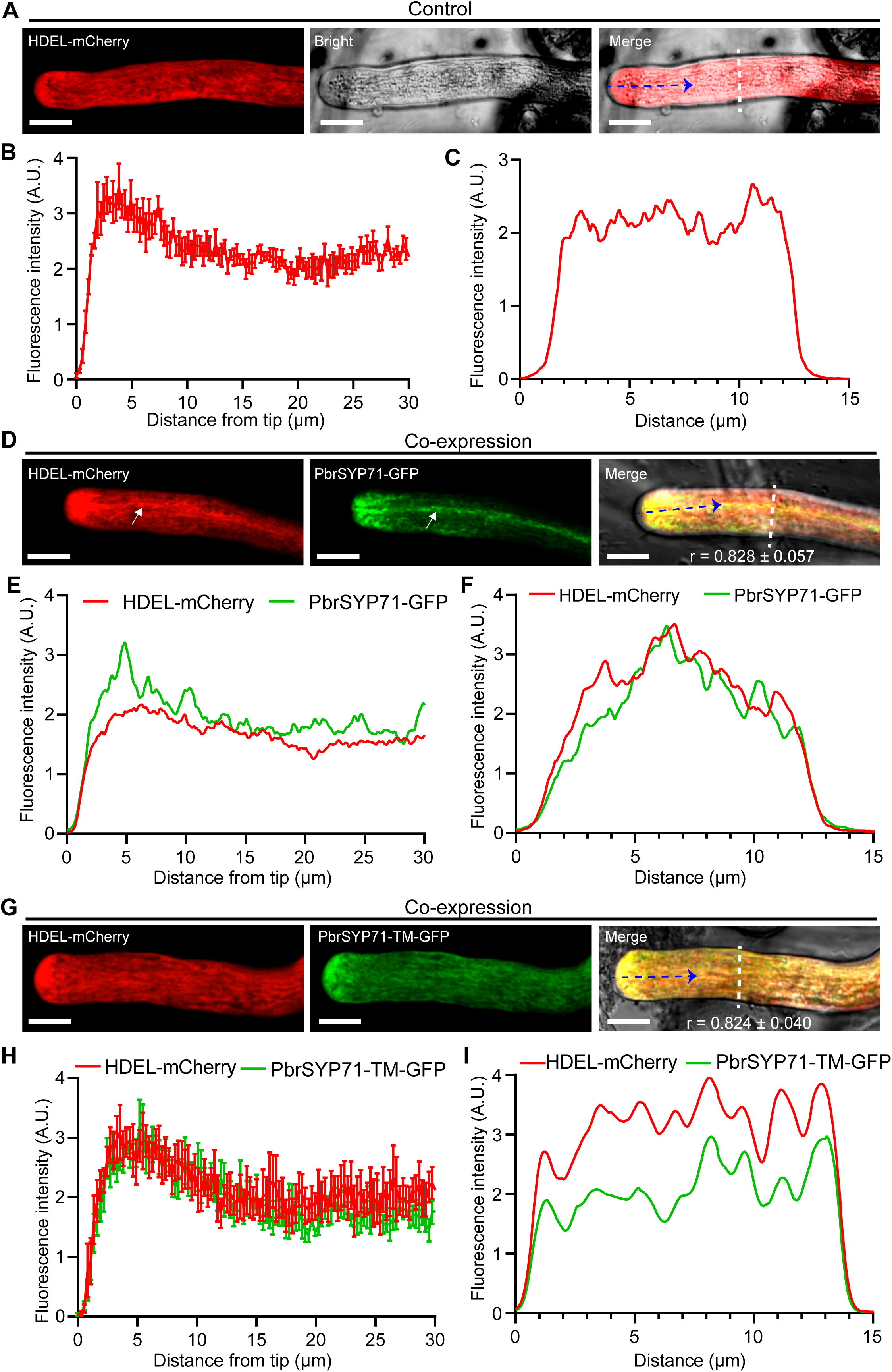
PbrSYP71 is localized to endoplasmic reticulum (ER) by transmembrane (TM) structure, and overexpression of *PbrSYP71* remodels the ER distribution in pear pollen tube. **A)** Image of pear pollen tube transiently expressing HDEL-mCherry. **B)** Fluorescence intensity of HDEL-mCherry at 30 µm distance from pollen tube tips along the blue dashed arrow in (A), n = 5 pollen tubes. **C)** Typical HDEL-mCherry distribution in the vertical growth direction of the shank of the pollen tube along the white dashed line in (A). **D)** Image of pear pollen tube transiently co-expressing PbrSYP71-GFP and HDEL-mCherry. Pearson correlation coefficients (r) indicate the extent of colocalization between PbrSYP71-GFP and ER-mCherry, data represent mean± SEM, n = 5 pollen tubes. **E)** Fluorescence intensity of HDEL-mCherry and PbrSYP71-GFP at 30 µm distance from pollen tube tips along the red thread in (D). **F)** Typical HDEL-mCherry and PbrSYP71-GFP distribution in the vertical growth direction of the shank of the pollen tube along the white dashed line in (D). **G)** Image of pear pollen tube transiently co-expressing PbrSYP71-TM-GFP and HDEL-mCherry. Pearson correlation coefficients (r) indicate the extent of colocalization between PbrSYP71-TM-GFP and HDEL-mCherry, data represent mean± SEM, n = 5 pollen tubes. **H)** Fluorescence intensity of PbrSYP71-TM-GFP and HDEL-mCherry at 30 µm distance from pollen tube tips along the red thread in (G), n = 5 pollen tubes. **I)** Typical HDEL-mCherry and PbrSYP71-TM-GFP distribution in the vertical growth direction of the shank of the pollen tube along the white dashed line in (G). For (A), (D), and (G), the exprssion of *HDEL-mCherry*, *PbrSYP71-GFP* and *PbrSYP71-TM-GFP* were driven by the NTP303 promoter. PbrSYP71-GFP and PbrSYP71-TM-GFP was transiently co-expressed with HDEL-mCherry in pear pollen tubes by particle bombardment. The pollen tube was cultured for 6 h after transformation. Scale bars = 10 μm.

In addition, the ER exhibited continuous movement towards the apex of the pollen tube, finally reversing direction in the apical region (*Movie supplement 3*). PbrSYP71-GFP and the HDEL-mCherry appeared clustered and moved towards the apex of the pollen tube and then reverse direction in the apical region (*Movie supplement 4*). Overexpression of PbrSYP71-TM-GFP did not induce abnormal distribution of HDEL-mCherry (*Movie supplement 5*).

We also verified that overexpression of *PbrSYP71* led to changes in the distribution of the HDEL-mCherry in tobacco leaf epidermal cells. The ER presented a typical network structure *(Figure* 3—*figure supplement 2B*). When *PbrSYP71* was overexpressed, the typical network structure of the ER disappeared and ER presented a clustered bundle structure (*Figure* 3—*figure supplement 2C*). PbrSYP71-GFP also colocalized with HDEL-mCherry with a high pearson correlation coefficient of 0.810± 0.011 (*Figure* 3—*figure supplement 2C, D*). The distribution of HDEL-mCherry showed no change due to overexpression of PbrSYP71-TM-GFP (*Figure* 3—*figure supplement 2E*). PbrSYP71-TM-GFP and HDEL-mCherry showed similar distributions, with a high pearson correlation coefficient of 0.767± 0.022 (*Figure 3* —*figure supplement 2E, F*).

In summary, PbrSYP71 was localized to the ER via TM structure and overexpression of *PbrSYP71* remodeled the ER distribution and led to accumulation distribution of ER. Therefore, we concluded that the overexpression of *PbrSYP71* influenced the distribution of ER in the pollen tube, ultimately leading to an inhibition of pollen tube growth.

### Knocking down the expression of *PbrSYP71* led to abnormal distribution of ER in pollen tube

To further verify the function of *PbrSYP71* in ER distribution, we performed ODN experiment to knock down the expression level of *PbrSYP71*. In control group (s-ODN), the red fluorescence of ER was continuously distributed throughout the cytoplasm of the pollen tube, with the highest fluorescence intensity at the sub apex of pollen tube (*Figure 4A, B* and *Movie supplement 6*). After treating the pollen tube with as-ODN-PbrSYP71, the ER distribution at the sub apex of pollen tube disappeared (*Figure 4A, B*), and the fluorescence intensity at the sub tip was significantly lower than that of the control group (*Figure 4C*), and the growth rate of pollen tubes significantly slowed down (*Movie supplement 7*). These results indicated that the uneven distribution of ER in the pollen tube was disrupted when the expression of *PbrSYP71* was suppressed.

**Figure 4.**
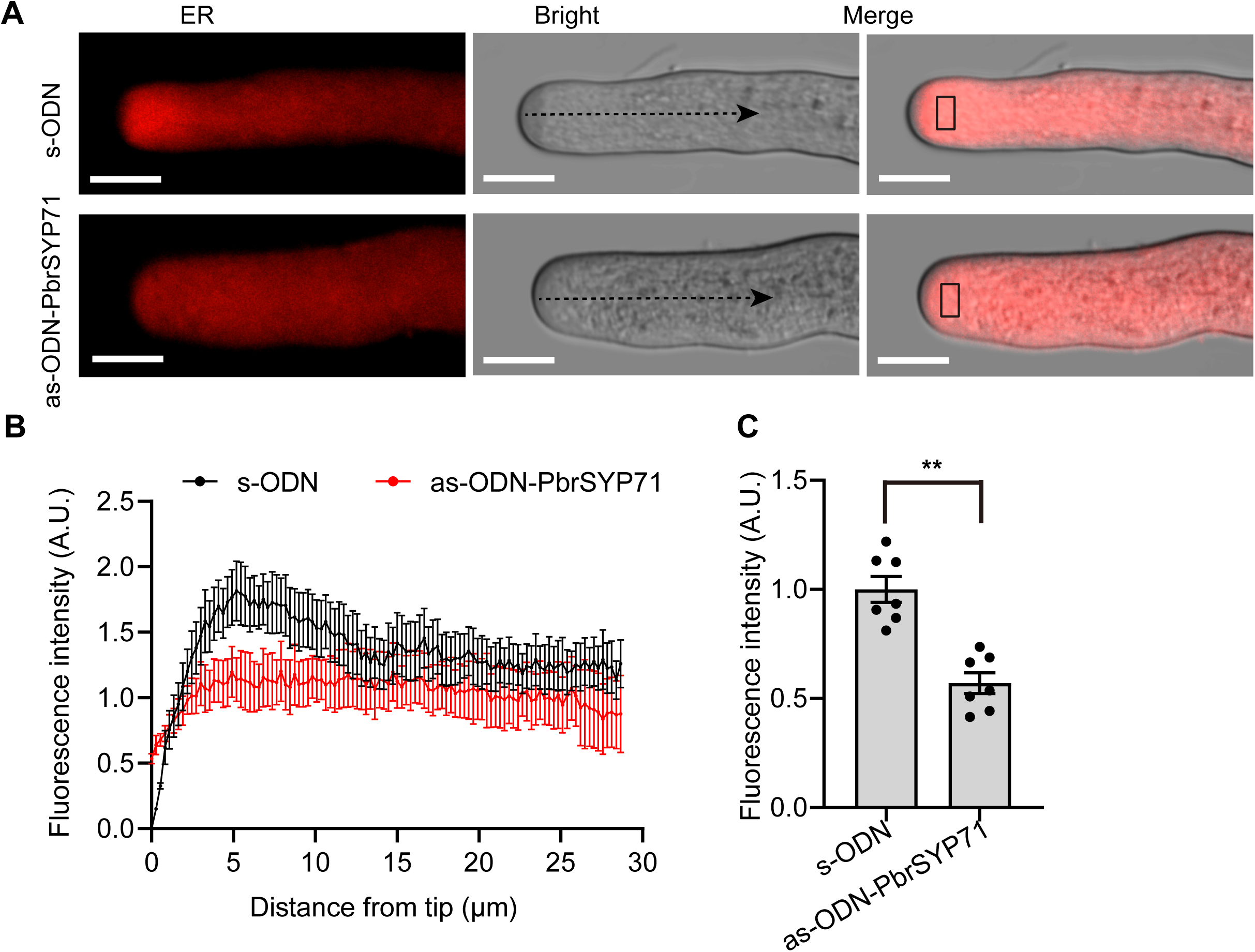
*PbrSYP71* is essential for endoplasmic reticulum (ER) distribution. **A)** The typical image of ER distribution of pear pollen tube treated with as-ODN-PbrSYP71 for 3 h. ER was stained by ER-tracker Red, s-ODN as control, scale bars = 10 μm. **B)** Fluorescence intensity of ER at 25 µm distance from pollen tube tips along the black thread in (A), n = 5 pollen tubes. **C)** Quantitative analysis of ER fluorescence intensity of pollen subapcial area at black box in (A), n = 7 pollen tubes. Data are mean± SEM, differences were identified using Student’s *t*-test, significant differences between means are indicated by ** (*p* < 0.01).

### ABD was the key domain for PbrSYP71 remodeling the ER distribution

Protein sequence conservation analysis indicates that PbrSYP71 contains an actin binding domain (ABD) at its N-terminus (*Figure 5A*). In order to investigate the function of ABD, we constructed a SYP71ΔABD-GFP vector without ABD structure and overexpressed it in pear pollen tubes (*Figure 5A*). When PbrSYP71ΔABD-GFP and HDEL-mCherry were co-expressed in pear pollen, the distribution and motion of HDEL-mCherry did not change, ER-mCherry was continuously distributed throughout the cytoplasm of the pollen tube, with the highest fluorescence intensity at the sub apex of pollen tube (*Figure 5B-D* and *Moive supplement 8*). PbrSYP71ΔABD-GFP and HDEL-mCherry showed similar distributions, with a high pearson correlation coefficient of 0.797± 0.053 (*Figure 5B*). Further research revealed that there was no significant difference in pear pollen tube length between the transient overexpression of PbrSYP71ΔABD-GFP and the overexpression of GFP (*Figure 5E, F*).

**Figure 5.**
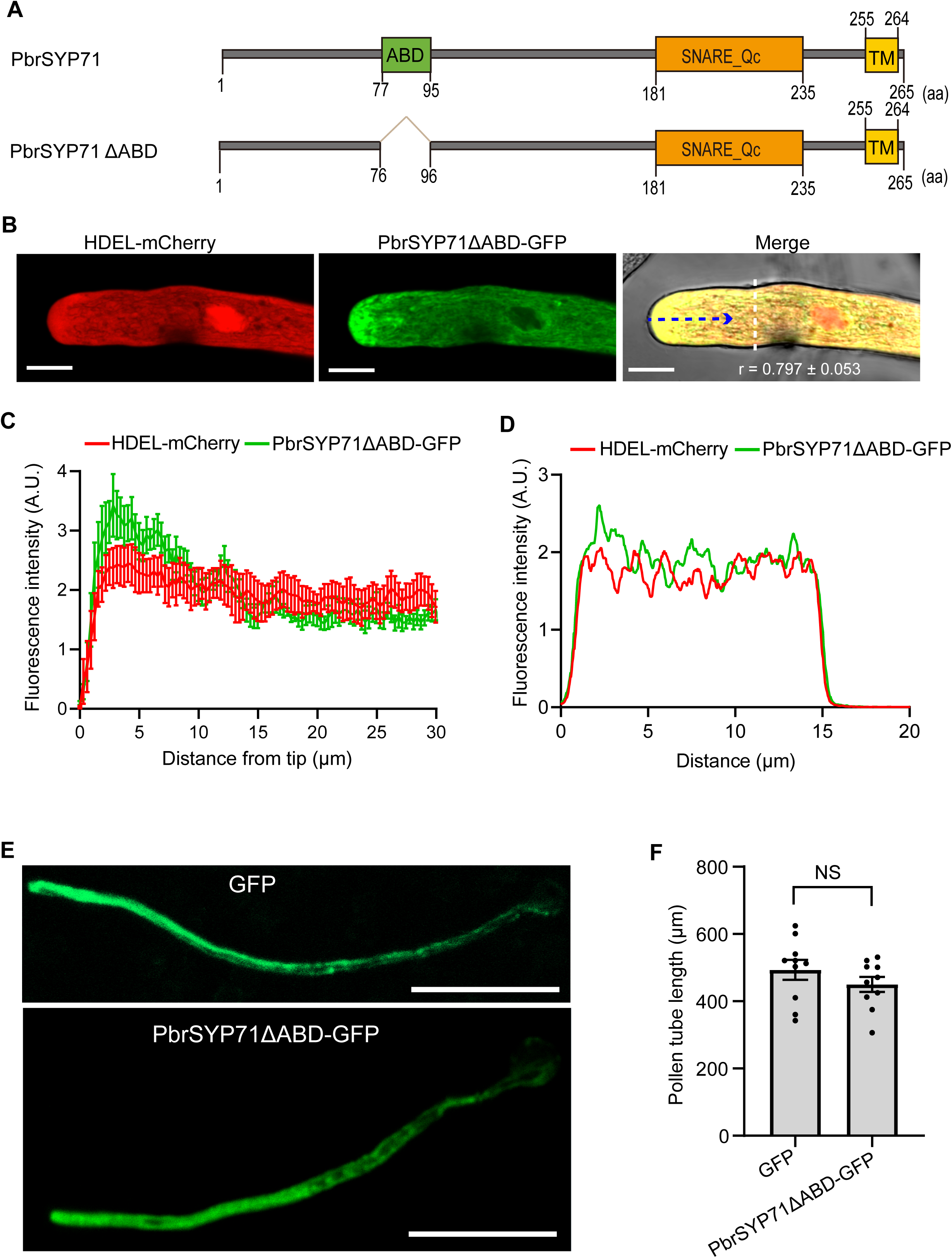
Actin binding domain (ABD) is key domain for PbrSYP71 remodeling the ER distribution and regulating pear pollen tube growth. **A)** Schematic diagram of PbrSYP71 and PbrSYP71ΔABD protein. ABD means actin binding domain, TM means transmembrane domain. **B)** Image of pear pollen tube transiently co-expressing PbrSYP71ΔABD-GFP and HDEL-mCherry. Pearson correlation coefficients (r) indicate the extent of colocalization between PbrSYP71ΔABD-GFP and ER-mCherry, data represent mean ± SEM, n = 5 pollen tubes. Scale bars = 10 μm. **C)** Fluorescence intensity of PbrSYP71ΔABD-GFP and HDEL-mCherry at 30 µm distance from pollen tube tips along the red thread in (B), n = 5 pollen tubes. **D)** Typical PbrSYP71ΔABD-GFP and HDEL-mCherry distribution in the vertical growth direction of the shank of the pollen tube along the white dashed line in (B). **E)** Image of pear pollen tube transiently expressing GFP or PbrSYP71ΔABD-GFP. The pollen tube was cultured for 6 h after transformation. Scale bars = 100 μm. **F)** Quantitative analysis of pollen tube length shown in (E), n = 10 pollen tubes. Data are mean ± SEM, differences were identified using Student’s *t*-test, no significant difference between means is indicated by NS (*p* > 0.05).

Meanwhile, we co-overexpressed PbrSYP71ΔABD-GFP and HDEL-mCherry in tobacco leaf epidermal cells. Both PbrSYP71ΔABD-GFP and HDEL-mCherry exhibited typical ER network structures (*Figure* 5—*figure supplement 1A*), and there was a high pearson correlation coefficients of 0.707± 0.033 (*Figure* 5— *figure supplement 1B*). Taken together, these results indicated that overexpression of ER-located PbrSYP71ΔABD-GFP did not alter the distribution of the ER in pollen tubes nor inhibit the pollen tube growth. Therefore, ABD was a key domain for PbrSYP71 remodeling the ER distribution and influences pollen tube growth.

### ABD was the key domain for PbrSYP71 to remodel ER to F-actin

First, we constructed LifeAct-mCherry for labeling the F-actin in pear pollen tubes. Then, we transiently co-overexpressed LifeAct-mCherry and PbrSYP71-GFP in pear pollen tubes. PbrSYP71-GFP and LifeAct-mCherry showed similar distributions, with a high pearson correlation coefficient of 0.719± 0.071 (*Figure 6B*). When PbrSYP71ΔABD-GFP and LifeAct-mCherry were co-expressed in pear pollen, PbrSYP71ΔABD-GFP and LifeAct-mCherry showed different distributions, with a lower pearson correlation coefficient of 0.528± 0.050 (*Figure 6C*). At the same time, we constructed the PbrSYP71-ABD-GFP vector for transient co-expression of PbrSYP71-ABD-GFP and LifeAct-mCherry in pear pollen tubes (*Figure 6A*). The results showed that PbrSYP71-ABD-GFP and LifeAct-mCherry exhibited similar distribution patterns, with a high pearson correlation coefficient of 0.815± 0.016 (*Figure 6D*). These results demonstrate that PbrSYP71 and PbrSYP71-ABD consistently co-localized with F-actin in pear pollen tube, highlighting the significance of the ABD domain as crucial for PbrSYP71 binding to F-actin.

**Figure 6.**
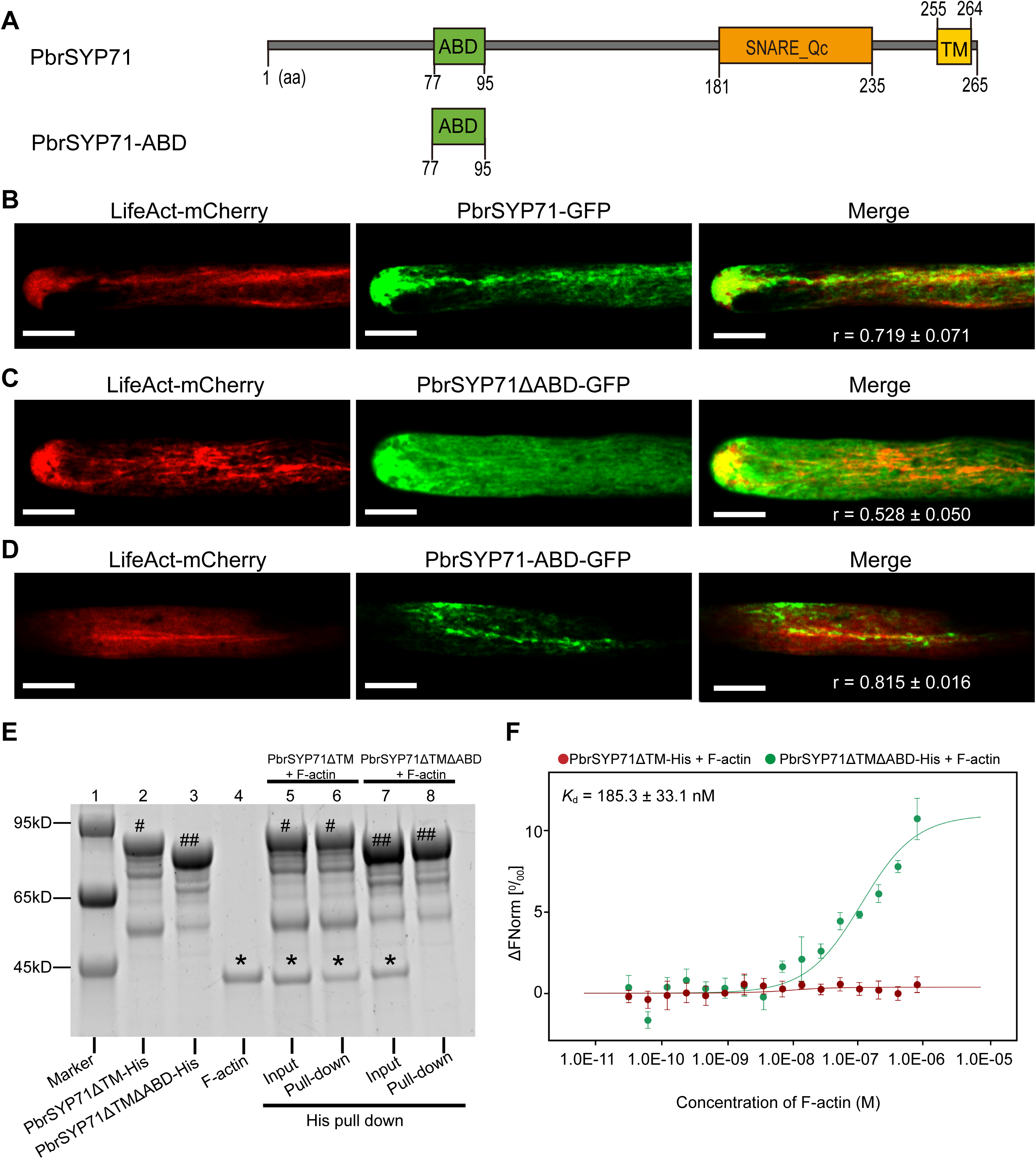
Actin binding domain (ABD) is the key domian for PbrSYP71 protein to remodel endoplasmic reticulum (ER) to F-actin and bind F-actin directly. A) Schematic diagram of PbrSYP71-ABD protein. **B)** The typical image of pear pollen tube transiently co-expressing LifeAct-mCherry and PbrSYP71-GFP. **C)** The typical image of pear pollen tube transiently co-expressing LifeAct-mCherry and PbrSYP71ΔABD-GFP. **D)** The typical image of pear pollen tube transiently co-expressing LifeAct-mCherry and PbrSYP71-ABD-GFP. **E)** ABD plays a crucial role in facilitating direct binding of the PbrSYP71ΔTM protein to F-actin. F-actin were detected after incubation with PbrSYP71ΔTM-His protein, F-actin were not detected after incubation with PbrSYP71ΔTMΔABD-His protein. PbrSYP71ΔTM-His, PbrSYP71ΔTM-His, and actin is marked by #, ##, *, respectively. For (B-D), Pearson correlation coefficients (r) indicate the extent of colocalization between PbrSYP71-GFP, PbrSYP71ΔABD-GFP, or PbrSYP71-ABD-GFP and LifeAct-mCherry, respectively. Data represent mean ± SEM, n = 5 pollen tubes.

The above results indicate that the ABD structure of PbrSYP71 is well colocalized with F-actin, suggesting that PbrSYP71 may bind to F-actin through the ABD structure. Therefore, we utilized the his pull-down method to verify whether PbrSYP71 binds to F-actin through ABD. Experimental results showed that PbrSYP71ΔTM-His (lacking TM domain of PbrSYP71) can bind to F-actin, while PbrSYP71ΔTMΔABD-His (lacking TM and ABD domain of PbrSYP71) cannot bind to F-actin (*Figure 6E*). In addition, microscale thermophoresis (MST) experiment results indicated that the binding affinity (*K*_d_) between F-actin and PbrSYP71ΔTM-His was 185.3± 33.1 nM, however, no binding affinity was detected between F-actin and PbrSYP71ΔTMΔABD-His (*Figure 6F*). These results indicated that PbrSYP71 attached to F-actin through the interaction between ABD domain and F-actin.

Due to the constant movement of the ER, to verify whether PbrSYP71 could provide power for the movement of the ER, we overexpressed PbrSYP71ΔTM-GFP, PbrSYP71-ABD-GFP, and PbrSYP71ΔABDΔTM-GFP in tobacco leaves, respectively. We observed that PbrSYP71ΔTM-GFP and PbrSYP71-ABD-GFP showed rapid movements (*Movie supplement 9* and *Movie supplement 10*), while the PbrSYP71ΔABDΔTM-GFP lacking ABD exhibited a loss of movement ability (*Movie supplement 11*). These results indicated that ABD was a key structure for PbrSYP71 to provide power for the movement of the ER.

## Discussion

The non-uniform distribution of ER in pollen tubes has been known for 17 years (Lovy-Wheeler et al., 2007; Hepler and Winship, 2015). However, the underlying molecular mechanisms remain largely unexplored. In this study, we discovered that a *SYP* gene, *PbrSYP71*, was specifically expressed in pear pollen tubes. Both overexpression and inhibition of *PbrSYP71* resulted in an anomalous ER distribution in the pollen tube and inhibited pear pollen tube growth. Further research found that ABD was the key domian for PbrSYP71 protein to remodel ER to F-actin and provided power for the movement of the ER.

### Syntaxin protein PbrSYP71 regulates cell tip growth

Syntaxin proteins are required for eukaryotes development as they play key roles in cell tip growth (Hong *et al*., 2010; Qi *et al*., 2016). Here, we identified a syntaxin protein – PbrSYP71 – that was involved in pear pollen tube tip growth. Not only did we demonstrate that *PbrSYP71* is essential for pollen tube growth, but we also found that overexpression of *PbrSYP71* severely inhibits pollen tube growth (*Figure 2*). Previously, syntaxin proteins are found to regulate cell tip growth in bacteria, fungi and model plant species *Arabidopsis*. For examples, in yeast, syntaxin Tlg1p protein mediates polarized growth by mediating delivery of chitin synthase III (Holthuis *et al*., 1998); in *N. crassa*, syntaxin NSYN1/2 proteins are involved in hyphal tip growth by regulating vesicle fusion (Gupta *et al*., 2003); in *Arabidopsis*, syntaxin of plant SYP121, SYP123, SYP132, and AtSYP81 proteins promote root tip growth by mediating vesicle fusion (Hirano *et al*., 2023; Wang *et al*., 2023), SYP124 is involved in pollen tube tip growth (Silva *et al*., 2010), SYP1s and SYP72 plays a key role in pollen development and germination (Li *et al*., 2019; Zhou *et al*., 2022). All these studies indicate that syntaxin protein is necessary for cell polarity growth, however, overexpression of the syntaxin gene has not been reported to inhibit cell polarity growth. Therefore, further elucidating the mechanism by which *PbrSYP71* inhibits pollen tube growth will contribute to a more comprehensive understanding of the function of syntaxin in cell development. To our knowledge, we are the first to identify the role of syntaxin protein in polarized cell growth in important crop plants (i.e. pear). All together, these results suggest that the function of syntaxin proteins in regulating cell tip growth is conserved across different species.

### PbrSYP71 regulates cell tip growth by affecting ER distribution

The distribution of ER is directly related to cell health and cell polarity (Zheng et al., 2022; Campanoni and Blatt, 2007). Our research showed that overexpression of *PbrSYP71* led to abnormal aggregation of the ER in the pollen tube, inhibiting pollen tube growth (*Figure 2; Figure 3*). When overexpression of *PbrSYP71*Δ*ABD*, which did not alter the distribution of the ER, pollen tube growth remained unaffected (*Figure 5*). Knocking down *PbrSYP71* resulted in the disappearance of uneven distribution of the ER, inhibiting pollen tube growth (*Figure 4*). This indicated that the uneven distribution of the ER was crucial for the polarized growth of pollen tubes. However, there have been few reports on the relationship between the distribution of endoplasmic reticulum, cell health and growth so far. In human U2OS cells, when ER distribution is disrupted, the distribution of other organelles is also affected, resulting in neurologic disorders and cancer (Zheng *et al*., 2022). ER accumulation at the root hair tip is necessary for the root hair development (Campanoni and Blatt, 2007). Here, we found that the distribution of the ER was closely related to the growth of pollen tubes, and abnormal distribution of the ER could lead to inhibition of pollen tube growth. These results provide new insights into exploring the relationship between ER distribution and plant cell polar growth, and serve as a reference for further research on the relationship between organelle distribution and cell growth.

In addition, PbrSYP71 protein, as t-SNARE protein, was localized at ER (*Figure 3*). Previously, syntaxin proteins are found to mediate vesicle fusion in together with ER. For example, ER-located t-SNARE proteins (i.e. syntaxin 18) mediate the fusion of COPI (coatomer complex I) vesicles, deriving by Golgi with ER and the fusion of legionella-containing vacuole with ER (Spang and Schekman, 1998; Hatsuzawa *et al*., 2006; Spang, 2013; Kawabata *et al*., 2021). Therefore, we propose that PbrSYP71 protein located at ER could mediate the fusion of vesicles with ER for maintaining the functioning of secretory system in pear pollen tubes. Previous studies on the function of syntaxin in cell polar growth mainly focus on regulating the fusion of vesicles and plasma membranes. For examples, syntaxin proteins located in plasma membrane, including syntaxin Tlg1p (in yeast), syntaxin NSYN1/2 (in *N. crassa*), SYP121 SYP123, SYP124, and SYP132 (in *Arabidopsis*), are involved in regulating the membrane fusion of secretory vesicle with plasma membrane for cell tip growth (Holthuis *et al*., 1998; Gupta *et al*., 2003; Silva *et al*., 2010; Hirano *et al*., 2023). Here, we showed that PbrSYP71 was located at ER and positively regulate vesicle secretion, which provided a specific target for studying the function of syntaxin in mediating vesicle fusion with ER during pollen tube growth.

In conclusion, syntaxin not only has the ability to regulate vesicle and target membrane fusion, but has also evolved to regulate the distribution of the ER. These results enrich the functionalities of syntaxin.

### ABD is a key domain of PbrSYP71 protein regulates the ER distribution

In animal and plant cells, some structural domain of the syntaxin protein plays a role in influencing protein distribution. For examples, in *Arabidopsis*, cytoplasmic domain of AtSYP73 protein interacts with actin, remodeling the ER to actin cytoskeleton (Cao *et al*., 2016). In INS-1 and CHO cells, H_abc_ domains of syntaxin 1A determine its cluster distribution without affecting its localization on plasma membrane (Yang *et al*., 2006; Fan *et al*., 2007). We found that ABD was the key domian for PbrSYP71 distribution and moving by anchoring to F-actin (*Figure 3; Figure 5; Figure 6; Movie supplement 10*). In conclusion, the cytoplasmic structure played a crucial role in influencing the distribution of syntaxin. Our results clarified the ABD of PbrSYP71 in regulating ER distribution, further broadening our understanding of the functions of syntaxin proteins in exocytosis process during polarized cell growth.

### The intracellular localization of PbrSYP71 protein is dependent on TM domain

Different syntaxin proteins have different subcellular localizations. For examples, syntaxin 1 and SYP131 (in mammals and *Arabidopsis*) are located on plasma membrane (Teng *et al*., 2001; Ichikawa *et al*., 2015); syntaxin 5 and SYP31 (in mammals and *Arabidopsis*) are presenting on the Golgi apparatus (Teng *et al*., 2001; Rui *et al*., 2021); syntaxin 18 and SYP73 (in mammals and *Arabidopsis*), as well as PbrSYP71 in pear found in our study, are located at ER (*Figure 3*; *Figure* 3—*figure supplement 2*) (Teng *et al*., 2001; Cao *et al*., 2016). The domains of syntaxin proteins that determine their specific subcellular localizations are also different for different syntaxin members (Kasai and Akagawa, 2001). We found that PbrSYP71 anchored the ER via TM domain (*Figure 3; Figure* 3—*figure supplement 2*). Additionally, the length of the TM domain determines the subcellular localization of syntaxins 3 (Watson and Pessin, 2001). The trans-Golgi network localization of syntaxin 6 depends on H2 domain and YGRL motif (Watson and Pessin, 2000), and the Golgi localization of syntaxin Sed5p relies on its the TMD amino acid, the COPI coatomer binding motif, and the folded-back conformation (Gao and Banfield, 2020). Thus, the conserved structures that influence the syntaxin subcellular localization differ between syntaxin proteins.

## Conclusion

Here, we presented a model to explain how *PbrSYP71* regulate pollen tube growth by regulating ER distribution in pear (*Figure 7*). PbrSYP71 protein is localized to ER via the TM domain and bridges ER with F-actin through ABD interaction with F-actin. This is the foundation of uneven distribution of ER with more accumulation at the sub-apical region in the pollen tube, facilitating the rapid growth of the pollen tube. When PbrSYP71 protein is insufficient, it can cause the disconnection between the ER and F-actin, leading to ER dispersion, reduced secretion vesicles, and ultimately inhibiting pollen tube growth. However, an excessive distribution of PbrSYP71 can cause the cluster distribution of the ER, also inhibiting pollen tube growth. In conclusion, we elucidate the key role of *PbrSYP71* in pollen tube growth by regulating ER distribution, providing new insights for the role of syntaxin proteins in regulation polarized cell growth.

**Figure 7.**
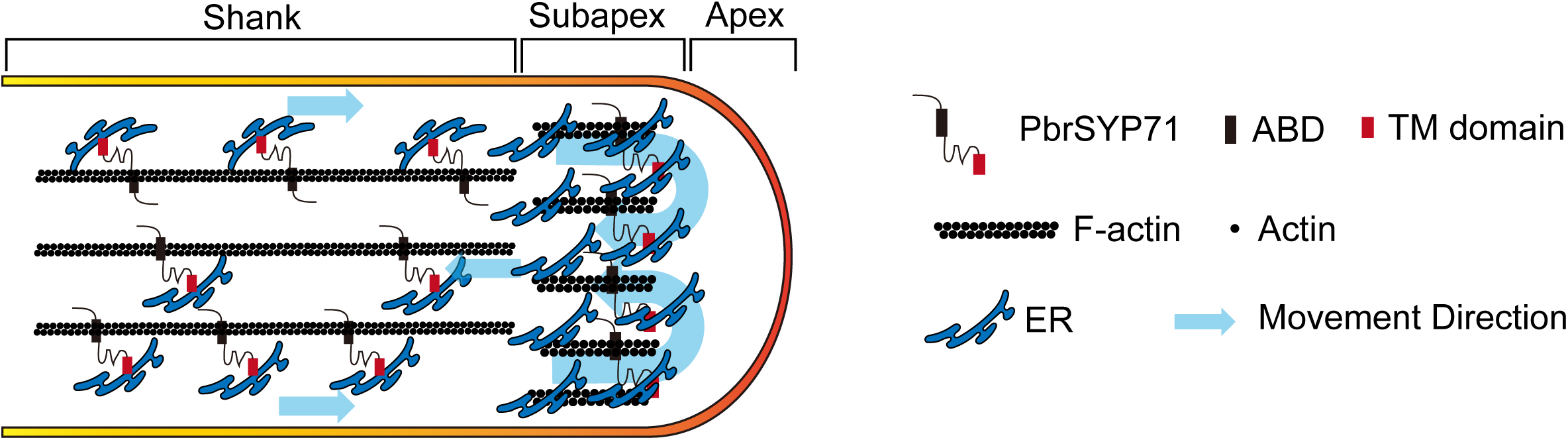
A working model for the role of PbrSYP71 in regulating pear pollen tube growth. Maintaining the suitable ER distribution is essential for the stability of pear pollen tube growth. PbrSYP71 is associated with the ER in pollen tube and bridges dynamically the ER membrane to the F-actin in virtue of the F-actin binding ability of ABD. The overexpression or absence of *PbrSYP71* disturbs the ER distribution, thereby inhibits pear pollen tube growth.

## Materials and methods

### Plant materials

Pear trees (*P. bretschneideri* cv. “Dangshansuli”) were grown in the Fruit Experimental Field of Nanjing Agricultural University, Nanjing, China (32°N, 118°E). Mature pollens from these trees were collected and stored at -20°C in a desiccator for later use.

### Identification and basic information analysis of *SYP* genes in pear

Protein sequences of 25 AtSYP were acquired from the TAIR (https://www.arabidopsis.org/) and these sequences were used as queries to perform BLASTP searches against the Pear Genome Database (http://pcwgdb.njau.edu.cn/) (Hu *et al*., 2023) by TBtools software (China) with an e-value cut-off of 1.0 × e^−5^ (Chen *et al*., 2023). Meanwhile, hidden Markov model profiles of the SYP domain (PF05739) were submitted to HMMsearch software to search for candidate SYP proteins in pear (Johnson *et al*., 2010). Redundant sequences from all obtained gene sequences were removed. SMART (http://smart.embl-heidelberg.de/) and the Pfam database (http://pfam.xfam.org/) were applied to detect SNARE functional domains in obtained sequences (Letunic *et al*., 2021; Mistry *et al*., 2021). The relative molecular weights and isoelectric points of PbrSYP proteins were determined by ExPASY (http://web.expasy.org/compute_pi/) (Artimo *et al*., 2012). The transmembrane structure and motifs of PbrSYP proteins were respectively predicted using CCTOP (https://cctop.ttk.hu/) (Dobson *et al*., 2015) and MEME (https://meme-suite.org/meme/) (Bailey *et al*., 2015). Actin-binding domain (ABD) LIG_Actin_WH2_2 was predicted by the Eukaryotic Linear Motif (ELM) program (Dinkel *et al*., 2016). The information of PbrSYP were given in Supplemental Table S1.

### Phylogenetic analysis

Phylogenetic analysis of 43 pear SYP members was performed by MEGA7.0 software (Kumar *et al*., 2016). Evolutionary analysis was conducted using 43 sequences of pear SYP proteins and 25 sequences of *Arabidopsis* SYP proteins. ClusterW alignment of all sequences was preferred, followed by the construction of a *SYP* evolutionary tree using the Neighbor Joining, with internal branches being taken 1000 times. Phylogenetic relationships of pear and *Arabidopsis* SYP7 subgroup proteins were conducted using the Neighbor Joining method in MEGA7.

### Pollen tube culture *in vitro*

The method of cultivating pollen *in vitro* can be found in our previous study (Tang *et al*., 2023). Mature pollen was taken out from -20°C and slowly thawed to room temperature, and evenly dispersed in a pollen tube medium (containing 0.5 mM calcium nitrate Ca(NO_3_)_2_, 1.5 mM boric acid (H_3_BO_3_), 25 mM 2-(N-morpholino) ethanesulfonic acid hydrate (MES) and 450 mM sucrose (sucrose) with pH 5.8, and cultured at 25°C/120 rpm. Then, the pollen tubes were collected by centrifugation at 2,000 rpm for 5 min, and then stored at -80°C.

### RNA extraction

Total RNA was extracted using a FastPure^®^ Universal Plant Total RNA Isolation Kit (Vazyme, China). The HiScript^®^ L1st Strand cDNA Synthesis Kit (Vazyme, China) was employed for reverse transcribing the RNA into cDNA.

### Quantitative real-time PCR (RT-qPCR) assays

The quantitative specific primers of *SYP* gene family members in pear were designed by the online Primer BLAST tool (https://www.ncbi.nlm.nih.gov/tools/primer-blast/) (Ye *et al*., 2012). LC480 II light cycler (Roche, Germany) instrument was used for qRT-PCR. The volume of each sample was 20 µL, which included 5 µL of specific primers with a concentration of 10 µM, 2 µL of cDNA with a concentration of 100ng/µL, 3 µL of ddH_2_O, and 10 µL of 2 × SYBR mixed dye (Vazyme, China). The reaction procedure for qPCR followed three steps, being that step 1 is incubating at 95L for 5 minutes, step 2 is incubating at 95 L for 3 seconds, followed by 60 L for 10 seconds and 72 L for 10 seconds, for a total of 45 cycles, and step 3 is drawing a dissolution curve to verify primer specificity. These data were analyzed using LightCycler480 software. *PbrUBQ* (Pbr035452.1) was used as an internal reference in pear pollen tube to normalize the expression levels of *SYP* genes. The relative expression level of *SYP* was calculated according to the 2^−ΔΔCt^ method. The primers of qRT-PCR assays were shown in Supplemental Table S3.

### Analysis of *PbrSYP* genes expression based on transcriptome data

The expression levels of *PbrSYP* genes were analyzed based on our previous transcriptomic data (http://pcwgdb.njau.edu.cn/) (Zhou *et al*., 2016; Hu *et al*., 2023) in 19 pear tissues including four seedling tissues (JSM, juvenile stem; CT, cotyledon; JLF, juvenile leaf; and RT, root), six flower organ tissues (OA, ovary; SP, sepal; ST, stamen; SY, style; PT, petal; and FLS, flower stalk), five fruit tissues (FTC, fruit core; FS, flesh ;FTS, fruit stalk; SD, seed; and PR, pericarp), flower bud (FLB), stem (SM), adult leaf (ALF), leaf bud (LFB). The transcripts per kilobase million (TPM) values were used for measuring the *PbrSYP* gene expression level at different developmental stages during pollen tube growth (MP, mature pollen; HP, hydrated pollen; PT, growing pollen tube; and SPT, stopped-growth pollen tube). The FPKM (reads per kilobase per million) values were used for measuring the gene expression level. These data of expression levels of *PbrSYP* genes were given in Supplemental Table S4, Table S5.

### Antisense oligodeoxynucleotide (as-ODN) assays

Antisense oligodeoxynucleotides (ODN) assays were performed as described previously in reference to the previous study (Chen *et al*., 2018). First, the mixture of ODN sequence and Lipofectamine 3000 (Thermo Fisher, USA) transfection reagent was incubated at room temperature for 15 minutes. Then the mixture was added to the pollen tube culture medium. The final concentration of ODN was 40 µM. Pollen was cultured for 3 h with 25L and 120 rpm. The images of pollen germination and pollen tube growth were obtained using an optical microscope, and the diameter and length of pollen tubes were measured using ImageJ software (https://imagej.nih.gov/ij/). The as-ODN Primer of PbrSYP71 gene was designed using online RNA fold website Server (http://rna.tbi.univie.ac.at//cgi-bin/RNAWebSuite/RNAfold.cgi). The sequence (AGGTCGCATCACAGCTAGGC) that does not exist in the pear genome as the s-ODN primer to treat the pollen tubes as the control group, and the specific complementary sequence (GCTTCTCGACGTCGTACTTG) predicted from the PbrSYP71 (Pbr025604.1) CDS as the as-ODN-PbrSYP71 primer to treat the pollen tubes as the treatment group. The primers of ODN were shown in Supplemental Table S3.

### Detection of cellulose in pollen tube cell wall

Cellulose staining of pollen tube cell wall was conducted based on our previous study (Li *et al*., 2022). First, prepared pollen tubes were washed with phosphate buffered saline (PBS) three times, and then fixed with 4% paraformaldehyde for 30 minutes. After fixation, pollen tubes were transferred to PBS containing 0.01% Pontamine Fast Scarlet 4B (S4B) (Sigma-Aldrich, USA), then incubated at room temperature for 5 minutes and washed with PBS three times. The dyed pollen tube was observed using a confocal microscope LSM800 (Zeiss, Germany). The fluorescence was measured using ZEN 3.1 software (Zeiss, Germany).

### Subcellular localization assays

The subcellular localization assays were performed based on sparkes’ method (Sparkes *et al*., 2006). Firstly, the CDS of *PbrSYP71*, *PbrSYP71-TM* (amino acid residue 236-265aa), and *PbrSYP71*Δ*ABD* without stop codon were cloned into the pCAMBIA1300-35S:GFP vector to construct 35S:PbrSYP71-GFP, 35S:PbrSYP71-TM-GFP and 35S:PbrSYP71ΔABD-GFP vectors. The CDS of HDEL was cloned into the pCAMBIA1300-35S:mCherry vector to construct 35S:HDEL-mCherry vector. We used HDEL-mCherry as a marker for the ER (Richard *et al*., 1992). Then, vectors were transformed into epidermal cells of *N. benthamiana*. Finally, the fluorescence was observed using an LSM800 laser scanning confocal microscope (Zeiss, Germany). The primers of subcellular localization assays were shown in Supplemental Table S3.

### ER labeling

ER-Tracker Red (Beyotime, shanghai, China) – a specific red fluorescent probe for ER – was used as a marker of the ER of eukaryotic cells (Wang *et al*., 2017; Su *et al*., 2021). ER of the pollen tube was labeled ER-Tracker Red as follows: pollen tubes were transferred to pollen culture medium containing 20 μΜ ER-Tracker Red and allowed to incubate for 30 min. The fluorescence of ER-Tracker Red was observed using an LSM900 laser scanning confocal microscope (Zeiss, Germany).

### Secretory vesicles labeling

Parton’s method was used to measure secretory vesicles (Parton *et al*., 2001). The pollen tubes were transferred to pollen culture medium containing 8 μΜ FM4-64 and allowed to incubate for 15 min. The fluorescence of FM4-64 was observed using an LSM900 laser scanning confocal microscope (Zeiss, Germany).

### Particle bombardment

Full-length cDNAs of *PbrSYP71*, *PbrSYP71-TM*, *PbrSYP71*Δ*ABD*, *PbrSYP71-ABD*, *HDEL*, and *LifeAct* were amplified using PCR and individually cloned into an overexpression vector driven by the NTP303 promoter (X69440.1) to generate PbrSYP71-GFP, PbrSYP71-TM-GFP, PbrSYP71ΔABD-GFP, PbrSYP71-ABD-GFP, HDEL-mCherry, and LifeAct-mCherry. HDEL-mCherry as ER marker, and LifeAct-mCherry as F-actin marker (Adriana and Benedikt, 2017). A mixture of 8.5 μL of gold particles (Bio-Rad, USA), 10 μL of 0.15 M spermidine, 2.5 μL of 2 μg/μL recombinant plasmid, and 29 μL of 2 M CaCl_2_ was prepared and vortexed continuously for 3 minutes. After vortexing, the gold microcarriers were subjected to centrifugation at 9,500 × g for 5 seconds, followed by removal of the supernatant. The gold particles were then washed with 200 μL of absolute ethanol, centrifuged again at 9,500 × g for 5 seconds, and eventually resuspended in 20 μL of absolute ethanol for the transformation process. The particle delivery system, PSD-1000/He, was set to the following parameters: 1,350 psi pressure, 20 mm Hg vacuum, 1-cm gap distance between the rupture disk and macrocarrier, and a 9-cm flight distance for the particles between the macrocarrier and the pollen samples. Prior to transformation, the pollen samples were positioned on solid germination medium containing 1.5% agarose. Fluorescence images were performed using a LSM800 laser scanning confocal microscope (Zeiss, Germany). The excitation wavelength and transmission range for emission were 488 nm/500 to 550 nm for GFP, and 561 nm/570 to 700 nm for mCherry. The detector gain was 700 V. The primers of particle bombardment were shown in Supplemental Table S3.

### Movie production

The video of PbrSYP71-GFP movement was produced using the following method. First, at least 20 consecutive fluorescence images were obtained using LSM900 laser scanning confocal microscope (Zeiss, Germany). Then, these images were played in chronological order (Frame Per Second=5)(Zeiss, Germany).

### Protein Expression and Purification

We used previous methods (Tang et al., 2023) to express and purify the His-tagged PbrSYP73ΔTM (lacking TM domain) and PbrSYP73ΔTMΔABD (lacking TM and ABD domain) recombinant protein. Full-length cDNAs of PbrSYP73ΔTM and PbrSYP73ΔTMΔABD were amplified by PCR and individually inserted into pCold-TF vectors. The pCold-TF vector functions as a fusion cold shock expression system that incorporates the trigger factor chaperone as a soluble tag. This trigger factor is a prokaryotic ribosome-associated chaperone protein with a molecular weight of 48 kDa, known to facilitate the cotranslational folding process of newly synthesized polypeptides. The primers are listed in Supplemental Table S3.

### His-tag pull down assay

To prepared 0.4 mg/ml F-actin working solution, 0.4 mg Pre-formed Actin Filaments (AKF99, Cytoskeleton Inc., >99% rabbit skeletal muscle actin) was dissolved in 1ml actin polymerization buffer, 10 mM Tris HCl, 2 mM MgCl_2_, 50 mM KCl, 1 mM ATP, 5 mM guanidine carbonate pH 7.5, (BSA02, Cytoskeleton Inc.,). To prepare the proteins for testing, the purified PbrSYP73ΔTM-His and PbrSYP73ΔTMΔABD-His were dialyzed in actin polymerization buffer.

Firstly, the tested proteins (40 µL, 1 mg/mL) were incubated with 40 µL 0.4 mg/ml F-actin working solution and actin polymerization buffer was added to reach a total volume of 100 µl. Set aside 20 μL of the mixture for use as the ‘input’. Then incubate this mixture at room temperature for 1 hour. Add 50 μL of Rigidarose Ni-NTA resin (His-tag) with a concentration of 5 μmol/ml and an average particle size of 75 μm. Incubate the mixture on a slow rotator for at least 2 hours at 4°C. Centrifuge the mixture at 800 × g and 4°C for 2 minutes. Discard the excess supernatant, leaving approximately 60 μL to prevent resin aspiration. Add 1 mL of actin polymerization buffer, gently resuspend the resin pellet, and centrifuge the mixture at 800 × g and 4°C for 2 minutes. Repeat the washing step at least three times. After the final washing step, ensure thorough removal of all residual liquid. Subsequently, elute the pull-down protein complex by boiling the resin in the SDS-PAGE protein loading buffer. Centrifuge the mixture at 12,000g for 1 minute and collect the supernatant as the ‘pull-down’. F-actin is boiled to depolymerize into actin, with a molecular weight of approximately 43 kD.

### Microscale thermophoresis (MST) assay

We used previous methods (Tang et al., 2023) to test the binding affinity (*K*_d_) between PbrSYP71ΔTM-His/PbrSYP71ΔTMΔABD-His and F-actin, respectively using the Monolish NT.115 instrument (Nano Temper Technologies, Germany).

### Statistical analysis

GraphPad Prism 8.2.1 (GraphPad Software, USA) was used to plot the results including their averages and the corresponding standard deviation. SPSS v22 (IBM, USA) was used to determine statistical significance. Significant statistical differences for two samples were analyzed using the Student’s *t*-test (*p* < 0.01). Significant differences for multiple samples were determined by ANOVA followed by Tukey’s multiple comparison test (*p* < 0.05).

## Author contributions

Juyou Wu, Mingliang Zhang, and Ming Qian designed the experiments; Mingliang Zhang, Chao Tang, Chi Lan, Dong Yue, Zhihua Xie, and Mengjun Sun conducted the experiments, with the help from Zongqi Liu, Zhu Xie, Hao Zhang, and Peng Wang; Mingliang Zhang, Juyou Wu, and Chao Tang wrote the manuscript; Ningyi Zhang, Peng Wang, Zhuqin Liu and Shaoling Zhang critically read and commented on the manuscript.

## Abbreviations list

ER: endoplasmic reticulum
SNARE: soluble N-ethylmaleimide-sensitive factor attachment protein receptor
F-actin: actin filaments
v-SNAREs: vesicle-associated SNAREs
t-SNAREs: target cell-associated
SYP: syntaxin of plants
s-ODN: sense oligodeoxynucleotide
as-ODN: antisense oligodeoxynucleotide
S4B: pontamine Fast Scarlet 4B
TM: transmembrane
aa: amino acids.

## Acknowledgements

We thank Dr. Barend H.J. de Graaf (Cardiff University) for offering us the vector NTP303:GFP. We thank Dr. Yuehua Ma from Central laboratory of College of Horticulture in Nanjing Agricultural University for her technical assistance in the use of laser confocal microscopy LSM800 (Zeiss, Germany). This work was supported by Jiangsu Agricultural Science and Technology Innovation Fund (CX(22)3044), the National Natural Science Foundation of China (32102358), the Natural Science Foundation of Jiangsu Province (BK20210394), the Project Funded by the Priority Academic Program Development of Jiangsu Higher Education Institutions, and the Earmarked Fund for China Agriculture Research System (CARS-28).

---

**Figure 1 — figure supplement 1. Neighbor-Joining phylogeny of *SYP* family proteins from *Pyrus* and *Arabidopsis*.** The phylogenetic tree was constructed by MEGA 7.0 software using full-length protein sequences. A bootstrap analysis was conducted with 1000 replicates. Different subgroups are distinguished by different color backgrounds (SYP1-SYP8), respectively. AtSYPs and PbrSYPs from *Arabidopsis* and *Pyrus*, respectively. The scale bar is 0.1.

**Figure 1—figure supplement 2. Conservative domains analysis of proteins of SYP7 subgroup. A)** Schematic diagram of SYP7 subgroup proteins demonstrating a SNARE-Qc domain. **B)** Predicted actin binding domains (ABD) and transmembrane (TM) domain have been identified in PbrSYP71-75.

**Figure 1—figure supplement 3. Conserved motif analysis of SYP proteins. A)** Phylogenetic tree of the *PbrSYP* and *AtSYP* famliy. **B)** Composition and position of 9 conserved motifs of PbrSYP proteins indicated by different color blocks. AtSYPs and PbrSYPs from *Arabidopsis* and *Pyrus*, respectively.

**Figure 1—figure supplement 4. Expression patterns of *PbrSYP* genes in different tissues of “Dangshansuli” pear.** Expression patterns of PbrSYP genes in 19 different tissues, including 4 seedling tissues (JSM, juvenile stem; CT, cotyledon; JLF, juvenile leaf; RT, root), flower bud (FLB), stem (SM), adult leaf (ALF), leaf bud (LFB), 6 flower organ tissues (OA, ovary; SP, sepal; ST, stamen; SY, style; PT, petal; FLS, flower stalk), 5 fruit tissues (FTC, fruit core; FS, flesh; FTS, fruit stalk; SD, seed; PR, pericarp). The expression levels were calculated as log2-transformed TPM values. white, yellow, and red indicate low, moderate, and high levels, respectively.

**Figure 1—figure supplement 5. The expression of *PbrSYP* genes during four development stages of pear pollen, including mature pollen (MP), hydrated pollen (HP), growing pollen tube (PT) and stopped-growth pollen tube (SPT).** Color scale represents log2-transformed FPKM values. The expression levels were calculated as log2-transformed FPKM values. white, yellow, and red indicate low, moderate, and high levels, respectively.

**Figure 2—figure supplement 1. The relative expression level of *PbrSYP73* and *PbrSYP75* was unaffected in pear pollen tube treated with as-ODN-PbrSYP71.** Data are mean± SEM, differences were identified using Student’s t-test, no significant difference between means is indicated by NS (*p* > 0.05).

**Figure 2—figure supplement 2. The cellulose content of the pollen tube was significantly decreased when the expression of *PbrSYP71* was knocked down. A)** Cellulose content of pollen tube wall was indicated using S4B dye under s-ODN and as-ODN treatments. Bars = 10 μm. **B)** Fluorescence intensity of S4B at 20 µm distance from pollen tube tips along the white thread in (A). n = 6. **C)** Quantitative analysis of the S4B fluorescence intensity at the apex of the pollen tube about 5 μm from the tip along black line in (A), at least 15 pollen tubes were measured. Data are mean ± SEM, differences were identified using Student’s *t*-test, significant differences between means are indicated by ** (*p* < 0.01).

**Figure 3—figure supplement 1. GFP and mCherry uniformly distribute throughout the cytoplasm of pollen tubes. A)** The image of pear pollen tube transiently co-expressing mCherry and GFP. The exprssion of mCherry and GFP were driven by the NTP303 promoter. GFP was transiently coexpressed with mCherry in pear pollen tubes by particle bombardment. The pollen tube was cultured for 6 h after transformation. Scale bars = 10 μm. **B)** Fluorescence intensity of GFP and mCherry at 30 µm distance from pollen tube tips along the red thread in (B), n = 3 pollen tubes.

**Figure 3—figure supplement 2. PbrSYP71 is localized to endoplasmic reticulum (ER) by transmembrane (TM) structure, and overexpression of PbrSYP71 remodels the ER distribution in tobacco leaf epidermal cells. A)** Schematic diagram of PbrSYP71 and PbrSYP71-TM protein. **B)** Image of tobacco leaf epidermal cells expressing the HDEL-mCherry (control). Scale bar = 10 μm. **C)** Image of tobacco leaf epidermal cells co-expressing PbrSYP71-GFP and HDEL-mCherry. **D)** Pearson correlation coefficients (r) indicate the extent of colocalization between PbrSYP71-GFP and HDEL-mCherry, data represent mean ± SEM, n = 3. **E)** Image of tobacco leaf epidermal cells co-expressing PbrSYP71-TM-GFP and HDEL-mCherry. F) Pearson correlation coefficients (r) indicate the extent of colocalization between PbrSYP71-TM-GFP and HDEL-mCherry, data represent mean ± SEM, n = 3.

**Figure 5—figure supplement 1. Actin binding domain (ABD) is key domain for PbrSYP71 remodeling the ER distribution. A)** Image of tobacco leaf epidermal cells co-expressing PbrSYP71-GFP and HDEL-mCherry. Scale bars = 5 μm. **B)** Pearson correlation coefficients (r) indicate the extent of colocalization between PbrSYP71ΔABD-GFP and HDEL-mCherry, data represent mean ± SEM, n = 3.

**Movie supplemental 1.** The movement of secretory vesicles during pear pollen tube growth.

**Movie supplemental 2.** The movement of secretory vesicles during pear pollen tube growth, when the expression of *PbrSYP71* was knocked down.

**Movie supplemental 3.** The movement of HDEL-mCherry in pear pollen tube.

**Movie supplemental 4.** The movement of HDEL-mCherry and PbrSYP71-GFP in pear pollen tube.

**Movie supplemental 5.** The movement of HDEL-mCherry and PbrSYP71-TM-GFP in pear pollen tube.

**Movie supplemental 6.** The movement of ER during pear pollen tube growth.

**Movie supplemental 7.** The movement of ER during pear pollen tube growth, when the expression of PbrSYP71 was knocked down.

**Movie supplemental 8.** The movement of HDEL-mCherry and PbrSYP71ΔABD-GFP in pear pollen tube.

**Movie supplemental 9.** The movement of PbrSYP71ΔTM-GFP in tobacco leaf epidermal cells.

**Movie supplemental 10.** The movement of PbrSYP71-ABD-GFP in tobacco leaf epidermal cells.

**Movie supplemental 11.** The movement of PbrSYP71ΔABDΔTM-GFP in tobacco leaf epidermal cells.

**Supplemental Table S1.** Detailed information on *SYP* gene family members in pear.

**Supplemental Table S2.** The 1D structure of PbrSYP proteins.

**Supplemental Table S3.** Primers used in this study.

**Supplemental Table S4.** Expression patterns of PbrSYP genes in different tissues of pear.

**Supplemental Table S5.** The expression of PbrSYP genes during four development stages of pear pollen tube.

